# Single molecule DNA methylation and hydroxymethylation reveal unique epigenetic identity profiles of T helper cells

**DOI:** 10.1101/2023.02.03.527091

**Authors:** Chloe Goldsmith, Olivier Fesneau, Valentin Thevin, Maria I. Matias, Julie Perrault, Ali Hani Abid, Naomi Taylor, Valérie Dardalhon, Julien C. Marie, Hector Hernandez-Vargas

## Abstract

Both identity and plasticity of CD4 T helper (Th) cells are regulated in part by epigenetic mechanisms. However, a method that reliably and readily profiles DNA base modifications is still needed to finely study Th cell differentiation. Cytosine methylation (5mC) and cytosine hydroxymethylation (5hmC) are DNA modifications that identify stable cell phenotypes but their potential to characterize intermediate cell transitions has not yet been evaluated. To assess transition states in Th cells, we developed a new method to profile Th cell identity using cas9-targeted single molecule nanopore sequencing and found that 5mC and 5hmC can be used as markers of cellular identity. Targeting as few as 10 selected genomic loci, we were able to distinguish major differentiated T cell subtypes as well as intermediate phenotypes by their native DNA 5mC/5hmC patterns. Moreover, by using off-target sequences we were able to infer transcription factor activities relevant to each cell subtype. Our analysis demonstrates the importance of epigenetic regulation by 5mC and 5hmC modifications in the establishment of Th cell identity. Furthermore, our data highlight the potential to exploit this immune profiling application to elucidate the pathogenic role of Th transition states in autoimmune diseases.

## Introduction

In general, adaptive immunity in both protective and autoimmune settings is orchestrated by the emergence of polarized effector CD4 T helper (Th) lymphocyte subsets of specialized function. Naive CD4 Th0 cells under the influence of cytokines and TCR signaling can give rise to different effector cells including Th1,Th2, Th17, T regulatory (Treg)^1,2^. Furthermore, plasticity is an important property of Th cells, whereby they adapt their functions in response to a changing environment forming a continuum of polarized phenotypes^3^. Hence, cells with intermediate Th1/Th17 and Th17/Treg phenotypes have been described in both mice and humans in association with autoimmune diseases and protective functions, respectively.

Both identity and plasticity of Th cells are regulated in part by epigenetic mechanisms^2,4^. Epigenetics refers to the heritable layer of information on top of a DNA sequence that regulates gene expression. The epigenetic landscape of cells makes them unique. Therefore, mapping epigenetic modifications represents an ideal way to discriminate cell types by their ‘epigenotype’^5,6^. DNA methylation is the most stable and most widely studied epigenetic mark and is involved in gene silencing^7^. DNA hydroxymethylation (5hmC) is relatively understudied, but recently has been linked to gene activity during differentiation ^8,9^. Because of their relative stability^10^ and cell-type specificity^8^, mapping 5mC/5hmC landscapes has the power to improve our understanding of Th cell identity and plasticity, and aid in the development of novel biomarkers for these cells.

Despite the evidence for 5mC/5hmC as a marker of cellular identity, in particular in Th cells, technical limitations have prevented this knowledge from reaching clinical use. Simultaneous detection of cytosine modifications (including 5mC and 5hmC) has been made recently available by the next generation of Nanopore long-read sequencers, which are able to provide such information in the native DNA context. Here, we take advantage of Nanopore sequencing to simultaneously profile 5mC and 5hmC in native DNA on naive Th cells polarized under Th0, Th1, Th2, Th17 and Treg conditions. In addition, given that Th17 cells differentiated in the absence of TGF-b are linked to experimental autoimmune encephalomyelitis (EAE) and autoimmunity when transferred into mice^11^, these cells were an appropriate model of pathogenic Th cells for the present study. Therefore, Naïve T cells were also polarized under Th17 conditions in the absence of cytokine TGF-β. This study opens new avenues towards the implementation of 5mC/5hmC in clinical immune profiling and identification of pathogenic immune cell subsets.

## Materials and methods

### Mice and mouse models

FoxP3-GFP reporter mice^12^ were housed in ventilated racks (with controlled temperature /hygrometry) in a pathogen-free facility at the Institut de Génétique Moléculaire de Montpellier. Animal care and experiments were approved by the local and national animal facility institutional review boards in accordance with French national and ARRIVE guidelines. All experiments were performed with adult (male) mice between the ages of 6-12 weeks of age.

### *In vitro* Th polarization

CD4^+^ T cells were isolated from male Foxp3-GFP reporter mice and naïve T cells were cell sorted and activated with anti-CD3/CD28 under different polarizing conditions: Th0 (neutral), Th1, Th2, Th17, Treg as well as Th17s differentiated without TGFβ (Th17-noTGFβ) representing an intermediate cell phenotype (**Supplementary tables 1 and 2**). Briefly, murine CD4^+^ T cells were purified using the MACS CD4^+^ T cell negative selection kit (Miltenyi Biotec) and naïve CD4^+^ T cells from WT, Foxp3-GFP were then sorted on the basis of a CD4^+^CD8^-^CD62L^+^CD44^-^GFP^-^CD25^-^ expression profile on a FACSAria flow cytometer (BD Biosciences). T cell activation was performed using plate-bound α-CD3 (clone 145-2C11, 1 μg/ml) and α-CD28 (clone PV-1, 1 μg/ml) monoclonal antibodies in RPMI 1640 medium (Life Technologies) supplemented with 10% FCS, 1% penicillin/streptomycin (Gibco-Life technologies) and - mercaptoethanol (50 μM). Cells were split 3 days later with medium supplemented with rhIL-2 (100U/mL). Cells were maintained in a standard tissue culture incubator containing atmospheric O2 and 5% CO2. 5mC raw data from Tregs included in this study was previously published by our group under the exact same experimental conditions^4^. These data were reanalyzed to compare an additional CD4 Th subset using updated basecalling algorithms and analysis pipelines.

### Flow cytometry and Cytometric Bead Array (CBA)

Immunophenotyping of cells was performed with fluorochrome-conjugated antibodies, and intracellular staining was performed after the fixation and permeabilization (ThermoFisher). Cells were labelled with fixable viability dye prior to fixation, followed by at least 3 hours of intracellular staining with the following anti-transcription factor antibodies; Foxp3-PeCy7, T-Bet-PE, Rorγt-BV650 and Gata3-APC. Cytokine production (IFN, IL-4, IL-17A, IL-10) was also assessed by Cytometric Bead Array (CBA) Kit (BD Biosciences) on supernatants collected at day 3 of polarization. Data analysis was performed using FlowJo (Tree Star software) and FCAPArray Software (CBA analysis).

### DNA extraction

DNA was extracted from 1×10^6 CD4 Th cells/ replicate using the Qiagen DNA mini kit according to manufacturer’s instructions (Qiagen Hilden, Germany). DNA was extracted from 10ng of mouse brain tissue as previously described^13^ using NEB Monarch High molecular weight DNA extraction kit according to manufacturer’s instructions (NEB Cat #T3050).

### Library prep and whole genome Nanopore sequencing

DNA was sheared using g-tubes (Covaris, USA) to ~10KB before library preparation with ligation sequencing kit (ONT Cat #SQK-LSK109) followed by whole genome sequencing on Nanopore MinION flow cells with pore 9.4.1 chemistry for 72h. Raw read quality was determined with pycoQC^14^.

### Cas9 enrichment, library prep and targeted Nanopore sequencing

A double cutting approach was used whereby 2x S. pyogenes Cas9 Alt-R™ trans-activating crispr RNAs (crRNAs) were designed for both upstream and downstream of each target loci as previously described^4,15^ (crRNA details and sequences provided in **Supplementary Table 3**). crRNAs were pooled in equimolar ratios (100 μM) and annealed to S. pyogenes Cas9 Alt-R™ tracrRNA (100 μM) (Integrated DNA Technologies, Iowa United States). crRNA•tracrRNA pool (10 μM) was incubated with Cut smart buffer (New England Biosciences Cat #B7204) and Alt-R^®^ S. pyogenes HiFi Cas9 nuclease V3 (62 μM) forming crRNA-tracrRNA-Cas9 ribonucleoprotein complexes (RNPs). 3000 ng of high molecular weight DNA was prepared by blocking available DNA ends with calf intestinal phosphatase (NEB Cat #M0525). DNA and dA-tails on all available DNA ends were cleaved by RNPs and Taq polymerase (New England Biosciences Cat #M0273) and dATP (New England Biosciences Cat #N0440), activating the Cas9 cut sites for ligation. Nanopore Native Barcodes (Oxford Nanopore Technology Cat #EXP-NBD104) were ligated to available DNA ends with NEB Blunt/TA Ligase Master Mix (New England Biosciences Cat #M0367) and up to 4 samples were pooled before ligation of Nanopore adapters with NEBNext^®^ Quick T4 DNA Ligase (New England Biosciences Cat #E6057) and loading onto a primed MinION flow cell (pore chemistry 9.4.1) and sequencing for 72h. Raw read quality was determined with pycoQC^14^.

### Detection of 5mC and 5hmC from Nanopore signal data

Raw fast5 files were basecalled with Megalodon (version 2.5.0, ONT) allowing the simultaneous delineation of 5mC and 5hmC by remora models (version 1.1.1, ONT) for the detection of modified bases.

### Detection of differential methylation

Data processing and statistical analyses were performed using R/Bioconductor (R version 4.0.3). Bed files containing modified cytosine information were transformed into a BSseq object for differential methylation analysis with dispersion shrinkage for sequencing data (DSS) as previously described^4,16^. DSS tests for differential methylation at single CpG-sites were evaluated using a Wald test on the coefficients of a beta-binomial regression of count data with an ‘arcsine’ link function. Differentially methylated regions (DMRs) were defined as those loci with at least 3 CpG sites within a distance of less than 50 bp, and with changes in > 50% of all CpG sites exhibiting p value < 0.05. Differentially methylated loci (DML) and DMR locations are available via GitHub (*https://github.com/hernandezvargash/Tcell.5mC.ID*). DMRs were plotted using the plotDMRs function of the dmrseq R package ^17,18^. Off-targets were analysed for transcription factor (TF) activity using the cistrome information associated with the MIRA R package^19^. After extracting all genomic binding regions corresponding to selected TFs, MIRA functions were used for aggregation of DNA methylation data and visualization of TF activity.

## Results

### Targeted Nanopore sequencing of *in vitro* polarized CD4 Th cells achieves high coverage of key loci

To study the relationship between DNA methylation and classic Th cell identity, polarized naïve Th0 cells towards various Th fates *in vitro*. Importantly, this approach does lead to a terminal differentiation state but rather a polarization towards different fate^20^. Th0 cells were isolated by FACs-sorting and activated under the following polarizing conditions towards the Th0, Th1, Th2, Th17 and Treg phenotypes using cytokine cocktails. And to test the sensitivity of this approach to detect potential pathogenic Th cell phenotypes, naïve T cells were polarized under Th17 conditions in the absence of cytokine TGF-β and are referred to as ‘Th17-noTGFβ’ cells (**Figure 1A**).

**Figure 1.**
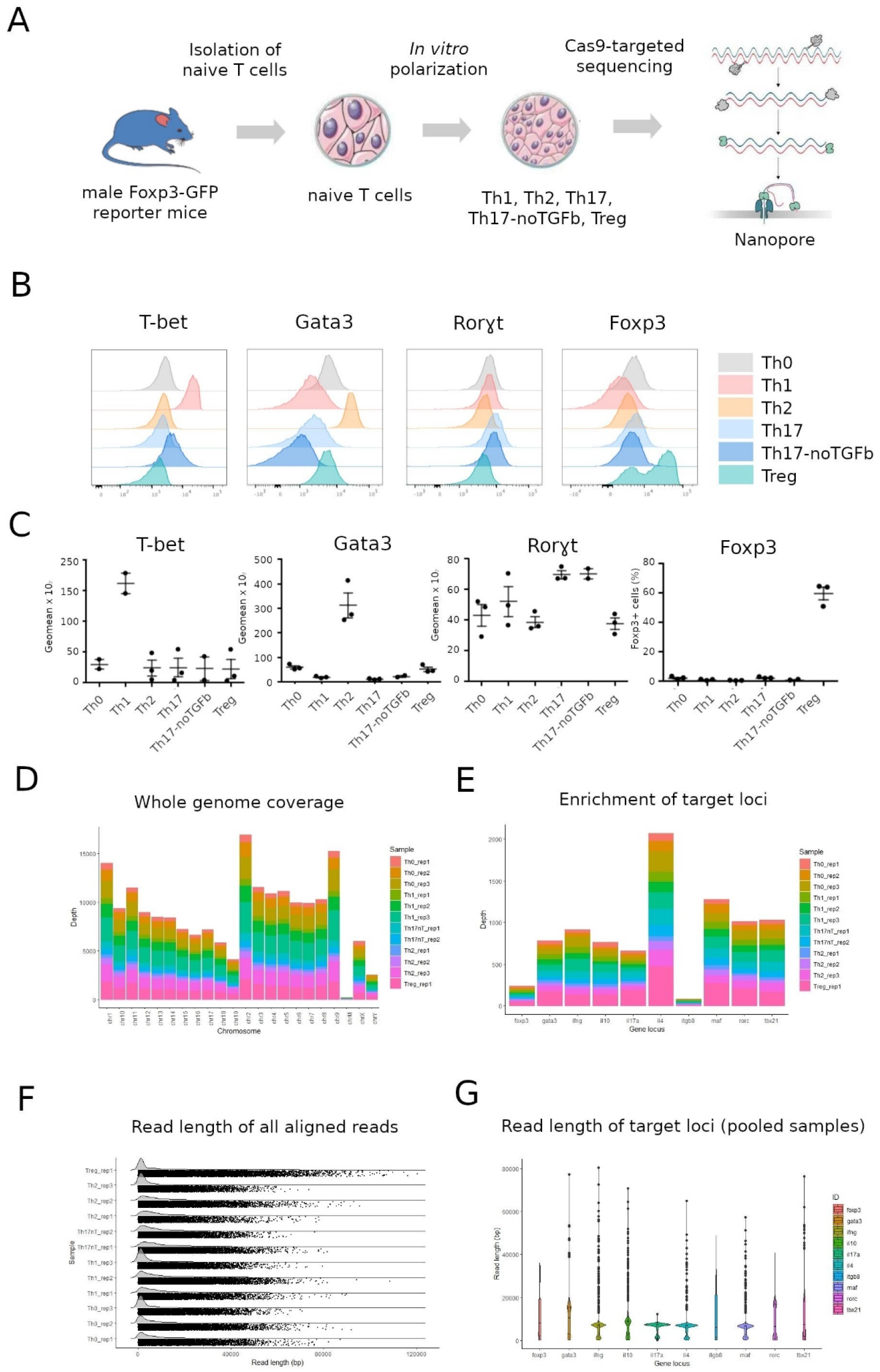
T-cell polarization, nanopore sequencing and validation. **A**. Experimental overview. T cells were *polarized in vitro* towards Th0, Th1, Th2, Th17, Th1/17 or Treg phenotype, followed by cas9 targeted sequencing with Nanopore. **B**. Transcription factor expression (ICS at day 4 of polarization). **C**. The pool of different experiments performed for transcription factor expression (ICS at day 4 of polarization). **D**. Nanopore sequencing yield and coverage. **E**. Length of all aligned reads in each sample, and length of reads aligning to targeted each region.

Specific transcription factor staining (**Figure 1B-C**) and cytokine production (**Supplementary Figure 1**) associated to Th cells were performed to estimate the efficacy of polarization for each condition. Expression patterns of transcription factors Tbet, GATA3 and Foxp3 appeared selectively upregulated in Th1, Th2 and Treg polarizing cells, respectively (**Figure 1B-C**) and correlated with the cytokine signature detected (IFNγ and IL-4 for Th1 and Th2 conditions respectively; **Supplementary Figure 1**). Expression of Rorγt was upregulated in Th17 and Th17-noTGFβ polarized cells, however the basal level of Rorγt for the other conditions (Th0, Th1, Th2 and Tregs) was higher than the other transcription factors (**Figure 1B-C**). Nevertheless, expression of Rorγt was consistently higher for all replicates of Th17 and Th17-noTGFβ conditions. MFI of Rorγt increased by 1.8 fold in Th17 and 1.7 fold in Th17-noTGFβ as compared to control conditions. Furthermore, only cells polarized towards a Th17 fate produce significant amounts of IL17 and did not produce IFNγ or IL4 at a detectable level (Figure 1C). Taken together, these data indicate an efficient polarization of Th17 and Th17-noTGFβ conditions.

To characterize the DNA methylation landscape of polarized Th cells, we developed a cas9 guided capture method coupled with Nanopore sequencing (**Figure 1A**). We identified important genes that could potentially be used to classify different Th subsets. This panel included DNA regions whereby differential epigenetic modifications in Th cell subsets have been previously reported^21–23^. In general, these genes include major transcription factors and cytokines specific to Th cell subsets, namely *Il17a, Maf, Rorc* and *Zfp362*^23^ (Th17); *Foxp3, Il10* and *Itgb8 (Treg); Gata3, Il4* and *Il10* (Th2); *Tbx21* and *Ifnγ* (Th1). A cas9 guided enrichment method was used to capture gene length regions prior to single molecule native DNA sequencing with Nanopore. This involved the design of 2 sgRNA’s upstream and downstream of each gene of interest employing a double cutting approach. It was then followed by synthesis of cas9 guided Ribonucleotide proteins (RNPs) to enrich for targets prior to Nanopore sequencing (**Figure 1D**) as previously described^4,6,15^. Up to 4 samples from each polarization assay were barcoded and multiplexed on each MinION flow cell. We profiled the 5mC and 5hmC landscape of whole gene length single DNA molecules in Th polarized subsets (Th0, Th1, Th2, Th17, Th17-noTGFβ and Tregs). In line with previous work on other cell types, the coverage of 10x was sufficient to detect a 25% change in DNA methylation^6^ and was therefore set as the minimum target coverage for this study. The typical sequencing yield achieved for each of these Nanopore runs was ~1GB (**Figure 1D-E**) which is in accordance with the literature^6,15^. Importantly, we exceeded the minimum coverage threshold of 10x for target regions (average coverage 45x, **Figure 1E**) and obtained full gene length reads (N50 = 21KB) on all targets (**Figure 1F-G**); this was sufficient for further processing to detect differentially methylated regions. These data highlight the efficacy of cas9 targeted sequencing for capturing important regions for Th cells.

### Classification of Th cell subsets using DNA methylation (5mC)

To basecall raw reads and detect modified bases, we used the latest high performing tool, Megalodon^15^ (https://github.com/nanoporetech/megalodon) combined with methylation models of the “Remora” repository. Megalodon is a command line tool that works by extracting high accuracy modified base calls from raw nanopore reads and anchoring the information rich basecalling neural network output to a reference genome (mm10). Global patterns of methylation were compared between different Nanopore sequencing runs and no differences were detected (**Supplementary figure 2A**). Conversely, there were significant differences between global 5mC distribution of CD4 Th cell subsets (**Supplementary figure 2C**); Th0 v Th2 (p<2.2e-16), Th0 v Th17 (p = 0.00086) and Th1 v Th2 (p<2.2e-16), Th1 v Th17 (p=0.0083) and Th17 and Th2 (<2.2e-16). Interestingly, the only cell subsets where no global difference was detected were Th0 v Th1 (p=0.45). These data could be explained in part by the genetic background of the mouse line which are known to be prone to develop certain Th cells. Nevertheless, these data highlight the potential specificity of methylation landscapes for different Th cell types.

Differentially methylated loci (DMLs) between each subset were determined, resulting in the identification of 573 significant (P<0.01) DMLs with greater than 10% difference (**Supplementary Table 4**). The number of distinct DMLs identified in Th1, Th2, Th17, and Treg cells when comparing to Th0 cells, were 59, 169, 103, and 79, respectively. Furthermore, a comparison of Th0 cells with Th1/17 cells identified 74 DMLs while 89 DMLs were identified between Th17 and Th17-noTGFβ cells, highlighting a potential effect of TGFβ alone on Th epigenetic identity.

Differentially methylated regions (DMRs) between subsets were determined by using DSS, resulting in 30 statistically significant (p<0.05) DMRs with a greater than 10% difference (**Figure 2**). In line with the selection of the chosen DNA regions, DMRs were largely negative compared to Th0 cells, highlighting that in these identity genes, 5mC decreased during T cell differentiation. Significantly, hierarchical clustering of DMRs was sufficient to cluster Th cells into their respective subsets (**Figure 2A**), with the one exception of the two Th17 samples differentiated in absence of TGF-β. Interestingly, plots of each DMR (**Figure 2B-F**) revealed the specificity of methylation profiles for a single cell type. Indeed, a DMR in the CpG island (CGI) shore located on *Tbx21* was unique to Th1 cells (**Figure 2B**), while the DMRs detected in *Il17a* were unique to Th17s (**Figure 2E)**. The same can also be said for DMRs detected in *Il4* and *Gata3* in Th2 cells (**Figure 2D**). Treg cells displayed a single DMR, hypomethylated relative to Th0 cells, in the region upstream of the *Foxp3* promoter (labeled according to the neighboring gene *Ppp1r3f*). Two DMRs were significantly hypomethylated in Th17 cells relative to naive Th0 cells, corresponding to the body of *Il1a7* and the *Rorc* promoter (**Figure 2E**). In contrast, the *Zfp263* locus was hypermethylated in these cells. These data highlight the power of characterizing DNA methylation profiling in a small subset of genes as means of determining Th cell identity.

**Figure 2.**
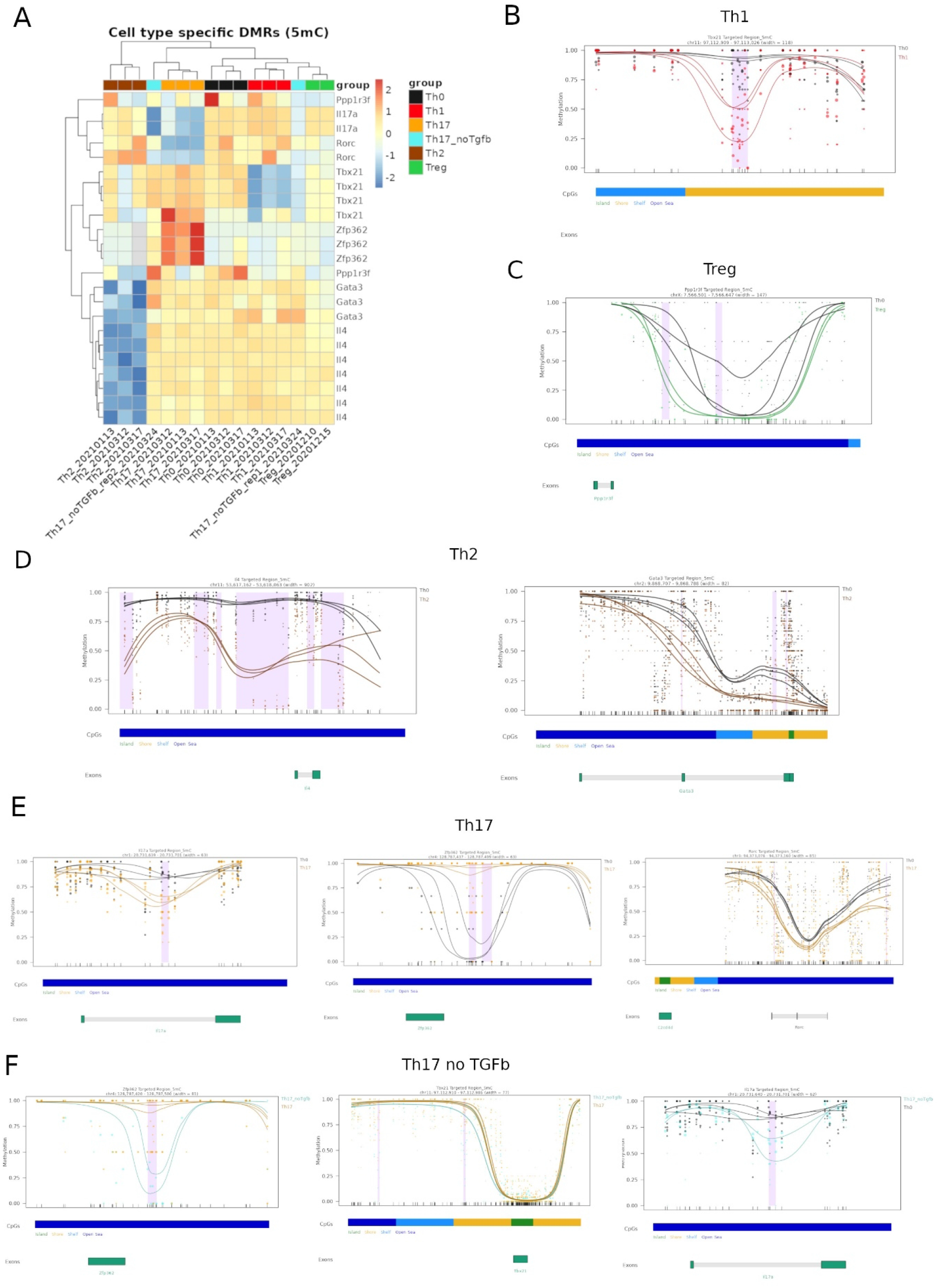
5mC landscape of immune identity genes in CD4 Th cells. **A.** Heatmap of cell type specific DMRs (5mC) in CD4 Th cells. **B**. Cell specific DMRs in Th1 v Th0 cells. **C**. Cell specific DMRs in Tregs v Th0 cells. **D**. Cell specific DMRs in Th2 cells. **E**. Cell specific DMRs in Th17 v Th0 cells. **F**. Cell specific DMRs in Th17 cells differentiated without TGFβ v Th17 cells. **Figure 3**. 5mhC landscape of immune identity genes in CD4 Th cells. **A**. Heatmap of cell type specific DMLs in CD4 Th cell subsets. **C**. DhMR identified at *IL4* locus in Th2 v Th0 cells.

Interestingly, although Th17-noTGFb cells exhibited demethylation the *Il17a* region to the same magnitude as Th17 cells differentiated in standard conditions (from 85% methylation in naive cells to ~50%; **Figure 2F**, right panel), *Zfp362* was not hypermethylated, remaining at similar levels as naive Th0 cells (**Figure 2F**, left panel). These cells also displayed less methylation at the *Tbx21* promoter, although less consistent across the two replicates (**Figure 2F**, middle panel). Thus, these data also suggest that 5mC methylation profiles can be used to evaluate the specific impact of a cytokine on Th differentiation.

### DNA hydroxymethylation (5hmC) is Th cell specific and indicative of gene activity

Though 5hmC is generally associated with increased gene expression when present in coding regions^16^, the relationship between 5mC and 5hmC in CD4 Th gene regulation is poorly described. This is of importance since 5mC and 5hmC have been shown as key players in tissue differentiation and cellular plasticity^8^. Therefore, we took advantage of Nanopore to simultaneously detect 5mC and 5hmC on the same native Nanopore reads, as described in Methods. While the detection of 5mC with Nanopore has been validated, the detection of 5hmC has not. Given that tissue derived from the central nervous system (CNS) is known for being high in DNA hydroxymethylation^24^, we used data obtained from DNA extracted from the brains of P13 OF1 mice as previously described^13^ as a positive control for 5hmC. We determined the global 5hmC levels in mouse brain and identified an enrichment of 5hmC in mouse forebrain enhancers in contrast to 5mC which was depleted in these regions (**Supplementary Figure 4**). These data demonstrated an efficient detection of 5hmC using Nanopore technology, and we proceeded to determine the 5hmC landscape of CD4 Th cell subsets.

Global patterns of 5hmC were compared between different Nanopore sequencing runs and no differences were detected (**Supplementary figure 2B**). Similarly to 5mC global distribution patterns, we identified significant differences between global 5hmC in Th cell subsets (**Supplementary figure 2D**). While no difference was detected between Th0 v Th1 (p=0.96), Th2 cells differed from other cell types Th0 v Th2 (p<2.2e-16), Th1 v Th2 (p<2.2e-16), and Th17 v Th2 (<2.2e-16). Th17 cells also differed from other cell types; Th0vTh17 (p=7.4e-09) and Th17vTh1 (p4.4e-09).

Differentially hydroxymethylated loci (DhMLs) between subsets were determined using DSS. These data revealed 100 significantly altered DhMLs (P<0.01; **Figure 3A**). A comparison of Th0 to Th cells identified DhMLs in specific subsets; 29 in Th1, 29 in Th2, 8 in Th17 and 14 in Tregs. A comparison of Th0 to Th17-noTGFβ identified 17 DhMLs, while a comparison of Th17 to Th17-noTGFβ identified 13 DhMLs.

**Figure 3.**
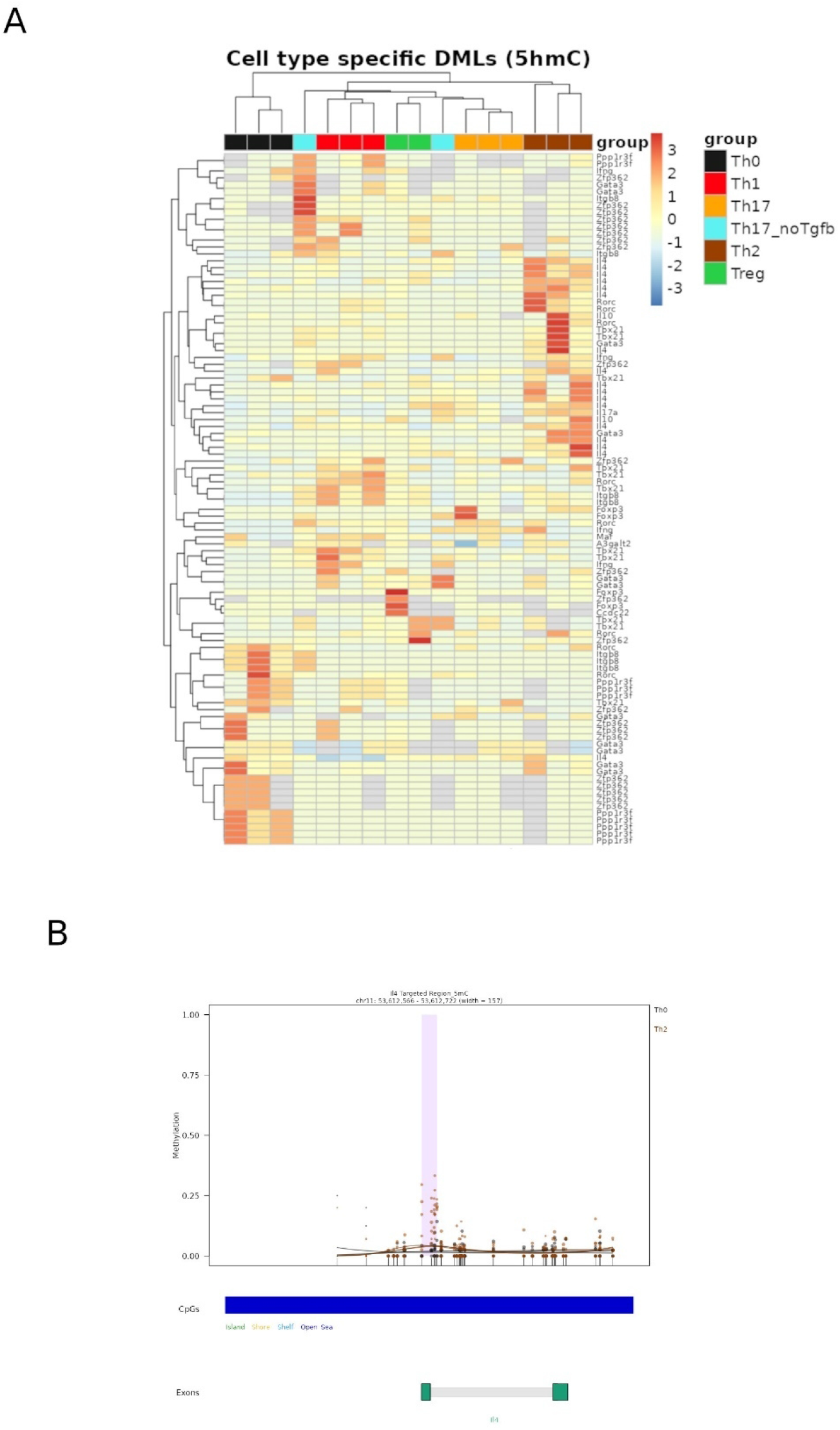
5mhC landscape of immune identity genes in CD4 Th cells. **A**. Heatmap of cell type specific DMLs in CD4 Th cell subsets. **C**. DhMR identified at *IL4* locus in Th2 v Th0 cells.

Differentially hydroxymethylated regions (DhMRs) were determined using DSS, as described in the methods. Only 1 DhMR was detected in *il4* comparing Th0 to Th2 cells. Interestingly, this DhMR also corresponded to an inverse DMR (DMR^-, DhMR^+) and was also associated with an increase in expression of IL-4 in Th2 cells (**Supplementary Figure 1**, **Supplementary Table 7**). In addition, we calculated the average 5mC and 5hmC for each target region (**Supplementary Figure 5**). As expected, 5hmC was largely concentrated in opposing target genes when compared to 5mC and was concentrated largely in predicted active genes in each subset (**Figure 3B**). Looking closely at the patters of 5mC and 5hmC in Th cells reveals the relationship between these modified bases in gene regulation, as well as the potential of 5hmC to serve as an efficient marker of active genes in Th cells.

### DNA methylation in transcription factor binding sites reveals their activity in Th cells

In addition to targeted regions, Nanopore sequencing captures around 1GB of ‘off’ target reads in each sequencing run. To strengthen the identification of Th cell subsets using DNA methylation, we determined the DNA methylation status of transcription factor (TF) binding sites in ‘on’ and ‘off’ target reads with Bioconductor package MIRA (Methylation-based inference of regulation activity). MIRA aggregates the DNA methylation in all known binding sites for chosen transcription factors. TF’s were chosen matching the known transcriptome orchestrators of Th cell subtypes *Tbx21 and Stat1* (Th1); *Gata3* (Th2), *Stat3* and *Runx3*(Th17), *Foxp3* (Treg)^25,26^. Importantly, *Bcl6* is a TF for T follicular helper cells^26^ and was used as a negative control for target Th cells. We aggregated 5mC located at known binding sites for the chosen TFs, representing hundreds of regions (**Figure 4**). In general, we observed a dip for all transcription factors, indicating hypomethylation, with the exception of the negative control TF *Bcl6*. In addition to plotting MIRA profiles, we determined the MIRA scores for each cell subset presented as boxplots (**Figure 4**). Validating the method, Tregs showed the highest MIRA scores for *Foxp3* binding sites, Th2 and Th17 displayed high scores for *Gata3* binding sites, and Th17 cells for *Runx3. Stat1* MIRA scores were highest in Th17 cells followed by Th2, with Th0 cells displaying a wide range in activity scores. *Stat3* MIRA scores were also highest in Th17 cells. Interestingly, for *Tbx21* binding sites the highest MIRA score was observed in Th17 cells followed by Th1 displaying a wide range in MIRA scores. Together, these data show that the methylation status of TF binding sites can be used as an indicator of the activity of Th cell subsets, further strengthening the relationship between 5mC and Th cell identity.

**Figure 4.**
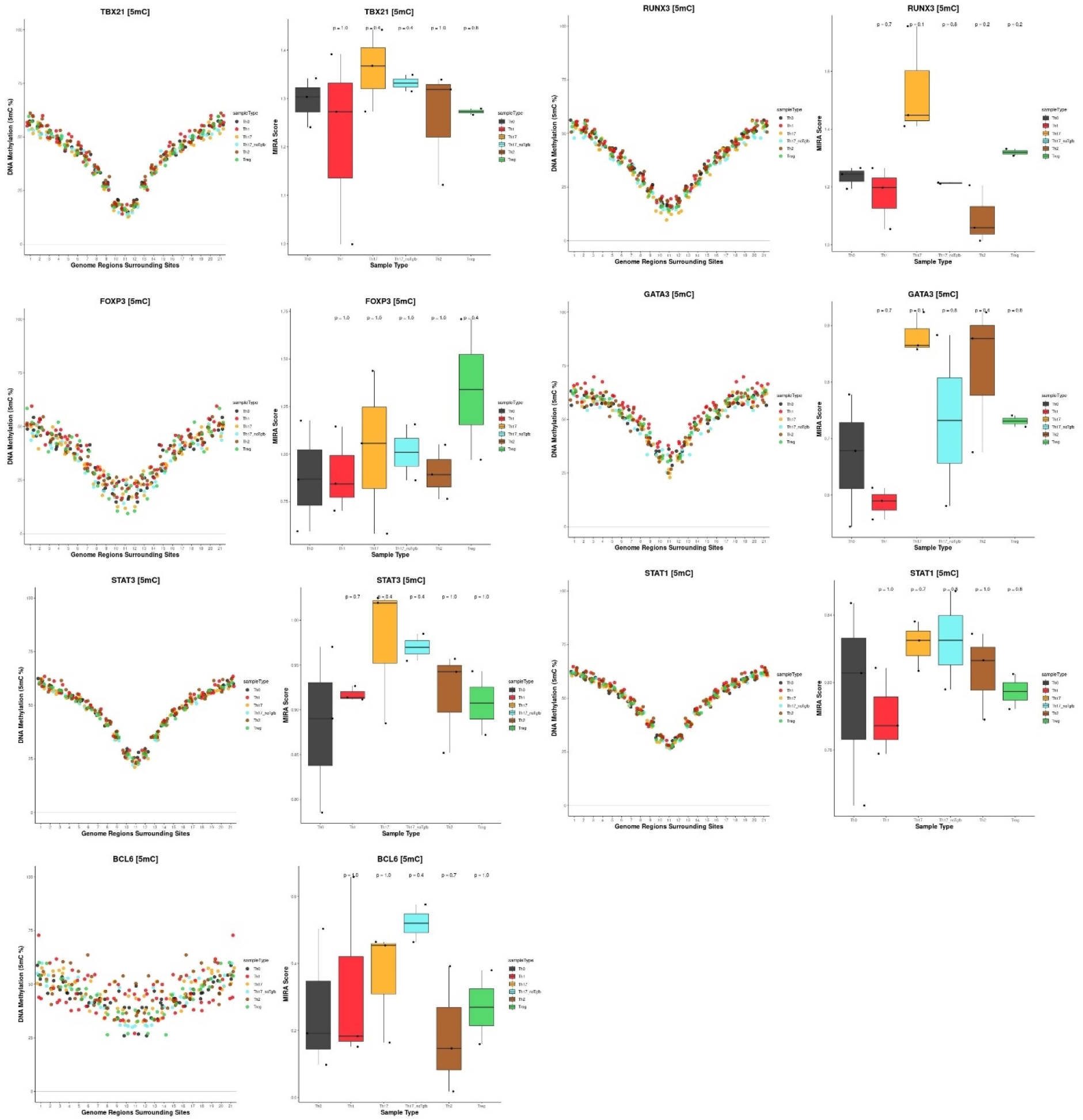
DNA methylation in Th cell transcription factor binding sites by MIRA. For each TF the MIRA profile is presented on the left and the aggregated score for each Th cell subset as boxplots on the right. p-values are based on comparison of each subset to the reference Th0 polarized cells

## Discussion

DNA cytosine modifications are known to be cell-type specific^1^, however the role of these modifications in Th cell identity is not well understood. The present study addressed the relationship between DNA methylation and Th cell identity, characterizing 5mC and 5hmC of murine CD4 Th cell subsets (Th0, Th1, Th2, Th17, Th17-noTGFβ and Tregs) by long read single molecule native DNA sequencing. We propose a novel approach able to identify differentially methylated and hydroxymethylated regions (DMRs and DhMRs, respectively) expected for each Th cell subset. Using this method, our data revealed that a combination of 5mC/5hmC profiles from different loci provides a map specific to each Th cell subtype.

We developed a cas9 targeted sequencing method to enrich important loci for Th cell identity. This method obtained high coverage of target genes compared to short read sequencing approaches, as well as whole gene length reads. Cas9 enrichment is an innovative technique that has been adopted to answer a variety of biological questions ranging from studying epigenetic heterogeneity in viruses^6^, important loci for cancer^15^ and neuromuscular disorders^27^. Regardless of the question, cas9 targeted enrichment coupled with Nanopore sequencing obtains anywhere from 0.5-2GB of data and 10 – 1000s of on target reads. In the present study, we observed variability between coverage of different target loci achieving the highest coverage of *Il4*, while the lowest with *Itgb8*. This coverage disparity is likely due to differences in size of target regions. *Il4* is a smaller gene and thus to obtain all regulatory information nearby we aimed to enrich in ~7Kb, while *Itgb8* is a larger gene and thus required enrichment of >100Kb. Shorter fragments elicit a higher coverage due to the nature of Nanopore’s preferentially sequencing shorter reads. Nevertheless, the on-target reads obtained in the present study fall well within that of the literature^15,27^.

Our data shows that 5mC and 5hmC in a selection of genes can accurately classify polarized Th cell subsets. Importantly, capturing long reads with high coverage of important loci revealed that DMRs were not only specific for a single subset; instead, some regions exhibited a range of methylation profiles. Other techniques relying on sodium bisulfite conversion are not able to distinguish 5mC from other modified bases such as 5hmC, which could lead to erroneous conclusions regarding the relationship between different modified bases and their role in gene regulation. Therefore, we focused our discussion on studies that consider both 5mC and 5hmC. Several studies have investigated the DNA methylation landscape of certain subsets (Th1 and Th2) using BS-seq and hydroxymethylation with immunoprecipitation techniques (hMeDIP)^28^. The authors highlighted, Th1 and Th2 cells could be distinguished by 5mC and 5hmC changes in important identity genes. We furthered this work by mapping 5mC and 5hmC in important loci for these and additional Th cell subsets (Th17, Th17-noTGFβ and Tregs) and identified unique 5mC patterns for each subset, and thus was able classify them by their epigenetic landscapes. Importantly, by characterizing methylation profiles in our identity gene panel, we were able to discriminate the unique Th17-noTGFβ cells (**Figure 2B**) indicating that an epigenetic approach has the potential to identify additional pathogenic and intermediate Th cell subsets. Emerging evidence suggests metabolomics and epigenetics work in concert to alter Th cell identity^1^. We have previously used this targeted sequencing approach to evaluate the relationship between metabolism and the epigenetic plasticity of regulatory T cells^4^. Here we found that in the presence of an abundance of certain metabolites (namely alpha-ketoglutarate), Tregs exhibited phenotypic plasticity adopting an inflammatory phenotype increasing expression of Rorc, Tbet and Ifnγ. These inflammatory cells were characterized by a specific epi-phenotype, with changes in DNA methylation correlating with gene expression aberration. These studies further support the utility of DNA methylation to identify plastic or pathogenic immune cell populations.

Indeed, DMR 5mC information from just a few loci can discriminate Th cell subtypes. We prepared spider plots to explore connections between DMRs in different genes and Th cell identity (**Supplementary Figure 6**). The different paths of Th polarization are illustrated by few dimensions in the 5mC data. Namely, by overlaying paths for Th17-noTGFb, Th1 and Th17s, we show that an aberrant path can be captured within these reduced dimensional frameworks. Moreover, as we learn from physiological and pathological Th differentiation, this strategy can be easily upgraded to cover a wider range of paths.

DNA hydroxymethylation is an emerging marker of cellular identity and has been studied largely in the context of cellular differentiation^9,29^. For detection of multiple modified bases (5mC, 5hmC etc), most techniques require samples to be split, and different modified bases to be detected separately. In the present study, we took advantage of Nanopore sequencing’s ability to determine the 5mC and 5hmC simultaneously. We found that global patterns of 5hmC in some T cell subsets were unique. Importantly, we also found that 5hmC across a selection of genes was able to accurately classify T cell subsets and was largely concentrated in ‘active’ genes. Our findings are in accordance with recent studies also highlighting that 5hmC in T cells is concentrated in the body of active genes^22,28^. However these studies indicated that terminally differentiated Th1 and Th2 cells have very little 5hmC^22^ compared to CD4^+^, CD8^+^ or DP T cells. This is not surprising considering the relationship between 5hmC and gene activity, and the high activity of genes during T cell differentiation^30^. ***Nevertheless, the present study is the first to showcase the specificity and often contrasting 5hmC patterns in a variety of differentiated CD4^+^ Th subsets. This is an important consideration for studies reporting molecular phenotypes of bulk PBMCs or whole blood^31^*.** Indeed, these findings also further support the role of 5hmC as a marker of gene activity and Th cell identity.

In addition to targeted DNA methylation analyses, we show here that off-target data can be used to effectively infer TF activity. As T cell identity is defined by the synchronized activity of TFs, we illustrate how this can be exploited to complement locus-specific results without additional extraction or sequencing workload. We foresee that a combination of both on-target and off-target data will be a more robust marker of Th cell identity.

An important consideration for interpretation of the present study; our samples were not purified after *in vitro* polarization and are known to be heterogeneous (cytokine treatment is not 100% efficient). However, we expect stronger discriminatory capacity in purified cell types. In addition, we selected important loci for Th cell subsets through literature review, which relied on known well annotated regions since there has yet to be a long read whole genome sequencing study of T cell subsets. Although we used analysis of off-target loci to support our main findings and develop an unbiased profile, there are potentially additional genomic regions with better discriminatory capacity that are yet to be identified. This is an important consideration for interpretation of our results.

In conclusion, gene DNA methylation modifications affect immunological pathways that may play an important role in disease. By mapping modified DNA bases (5mC and 5hmC) at single molecule resolution we show that specific epigenetic footprints are able to distinguish some of the major groups of *in vitro* polarized T cells (i.e. Th0, Th1, Th2, Tregs and Th17s). Therefore, we show that DNA methylation and hydroxymethylation are cell specific and have the power to identify and classify Th cell subsets and indicate their presence in heterogenous systems. This is an important consideration for bulk epigenome sequencing studies of heterogenous tissue like whole blood, PBMCs or tumor tissue. While, pathogenic phenotypes can also be identified by 5mC/5hmC (Th17-noTGFb), further studies are required to determine if this method has clinical applicability. Importantly, additional studies are needed to characterize the epi-phenotype of human and disease specific T cells.

## Author Contributions

CG conducted all sequencing experiments.

MM and JP performed all *in vitro* differentiation experiments.

JM, OF and VT participated in study design.

CG, AH and HH performed bioinformatic analyses

CG and HH wrote the manuscript.

VD, NT and HH supervised the work.

HH conceived the study.

All authors discussed the results and manuscript text.

## Data availability

All sequencing data was made available by depositing to XX. Ascension number XX.

Bioinformatics code: https://github.com/hernandezvargash/Tcell.5mC.ID.

A shiny application accompanies this manuscript: http://20.56.136.251:3838/test.app/.

## Funding

This work was supported by the Institut Nationale de la Sante et de la Recherche Medicale (INSERM) Plan Cancer AAP “Single Cell”, édition 2019, No. 19CS144-00 (project: *Molecular and cellular map all along anorectal cancer development);* the Institut National du Cancer AAP PreNeo 2019 (project: *Etude des mécanismes de développement des cancers anorectaux dans un modèle de néoplasie intra-épithéliale);* the Fondation MSD Avenir, programme ERICAN 2020 (project: *Deciphering the epigenetic mechanisms responsible of the Th17 plasticity associated with tumor development);* the Agence Nationale de la Recherche (ANR) 2020, Allocation No. RPV20018CCA (project: *Etude de la plasticité et du potentiel de différentiation des cellules des zones de transition pendant l’homéostasie normale et perturbée*); and the Fonds Amgen France pour la Science et l’Humain 2021 (project: *Establishing a clinically committed pipeline to identify T cell and tumor epigenotypes associated with cancer and its response to immunotherapy*).

## Competing interests

CG and HH have received travel and accommodation support to attend conferences for Oxford Nanopore Technology. The authors declare no additional competing interests.

## Notes

https://github.com/hernandezvargash/Tcell.5mC.ID

http://20.56.136.251:3838/test.app/

