## Supplementary Figures and Tables 1-3 for "Single molecule DNA methylation and hydroxymethylation reveal unique epigenetic identity profiles of T helper cells"

### Supplementary figure and table legends

**Supplementary Figure 1**. Cytokine expression of Th cell subsets; Th0, Th1, Th2, Th17 and Tregs, Th17-TGFb indicate Th17 cells differentiated without TGFb.

**Supplementary Figure 2.** Global **5mC** and **5hmC** landscape CD4 Th cells. **A**. Distribution of 5mC in different Nanopore runs. **B**. Distribution of 5hmC in different Nanopore runs. **C**. Distribution of 5mC and genomic features in different polarized Th subsets. **D**. Distribution of 5hmC and genomic features in different polarized Th subsets.

**Supplementary Figure 3.** Raw read accuracy of each Nanopore run determined by pycoQC

**Supplementary Figure 4**. DNA hydroxymethylation landscape of mouse brain tissue. **A**. Distribution of 5mC and 5hmC in mouse brain tissue. **B**. 5mC and 5hmC levels detected within 5KB of CpG islands (CGIs). **C**. 5mC and 5hmC levels detected within 5KB of gene bodies. **D**. 5mC and 5hmC levels detected within 5KB of Transcription start sites (TSS). **E**. 5mC and 5hmC levels detected within 5KB of forebrain enhancers. **F**. 5mC and 5hmC levels detected within 5KB of adipose tissue enhancers.

**Supplementary Figure 5**. DNA methylation and hydroxymethylation across whole target genes. Z-scores of mean 5mC and 5hmC calculated across all target regions.

**Supplementary Figure 6**. Relationship between 5mC in identity genes and Th cell identity. Spider plots displaying the methylation (5mC) status of the largest DMR for each significant gene indicated in Th cell subsets. **A.** Each spider plot represents a different replicate for Treg, Th0 and Th2s. **B.** Spider plots overlay of a representative replicate for Th0, Th2 and Tregs. **C.** Each spider plot represents a different replicate for Th17-noTGFb, Th1 and Th17s. **D**. Spider plots overlay of a representative replicate for Th17-noTGFb, Th1 and Th17s.

**Supplementary Table 1.** Key resources table; Antibodies, recombinant proteins and Flow cytometry reagents.

**Supplementary Table 2**. T cell polarizing conditions used for each Th subset.

**Supplementary Table 3**. sgRNA designed using IDT design tool for immune identity gene panel.

**Supplementary Table 4.** DMLs

**Supplementary Table 5.** DMRs

**Supplementary Table 6.** DhMLs

**Supplementary Table 7.** DhMRs

### Supplementary Figures

A

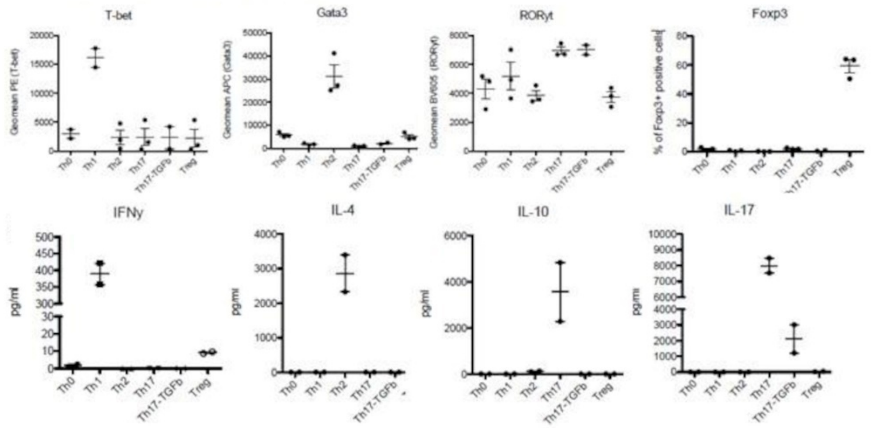


**Supplementary Figure 1.**

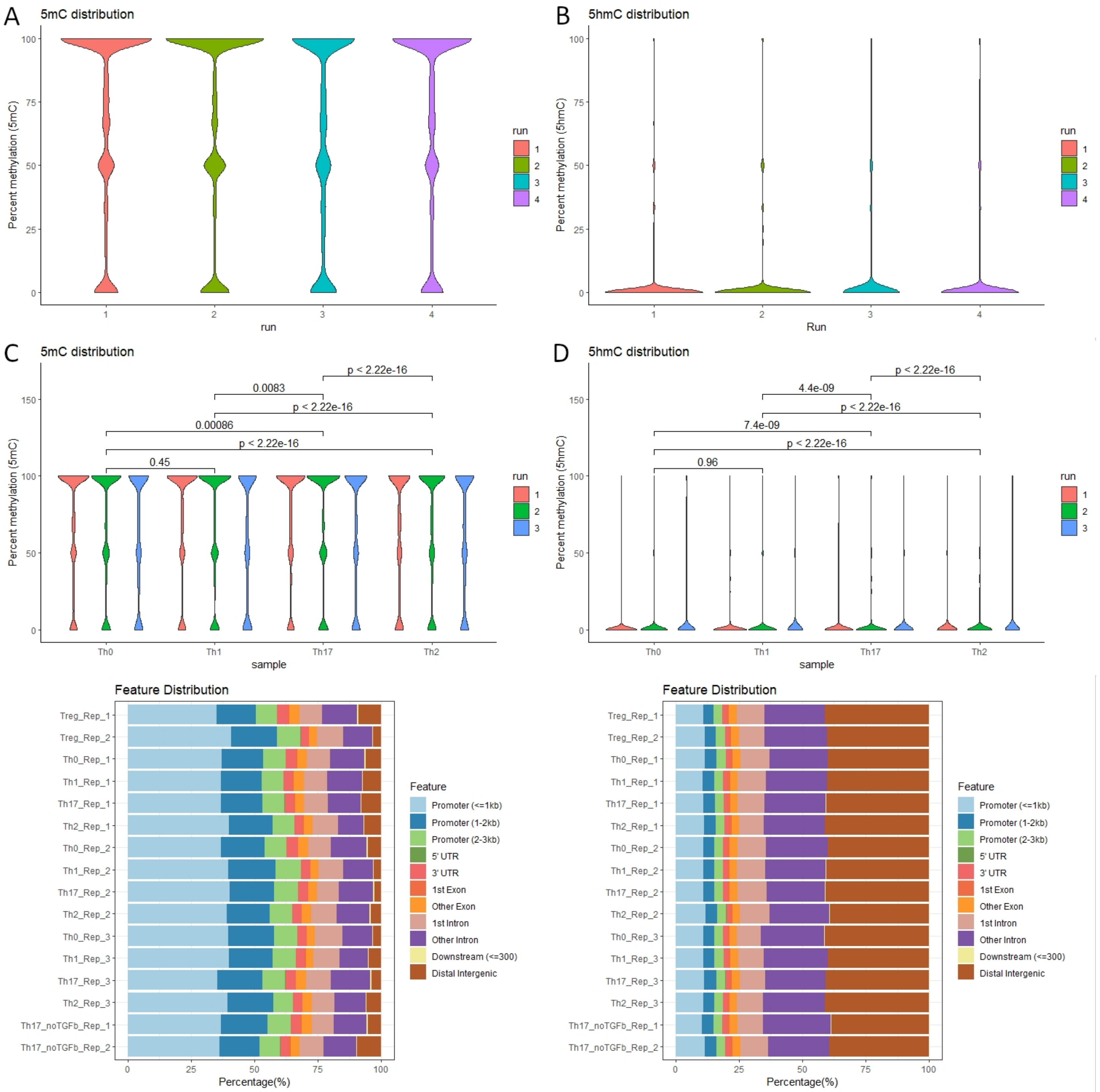


**Supplementary Figure 2.**

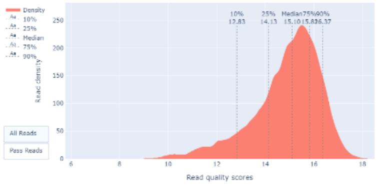


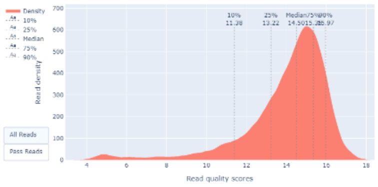


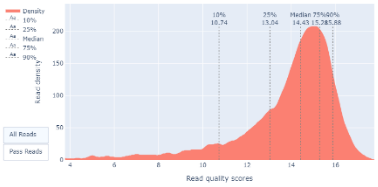


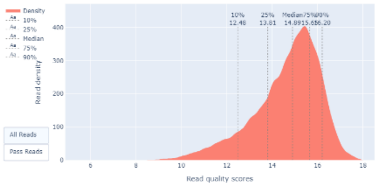


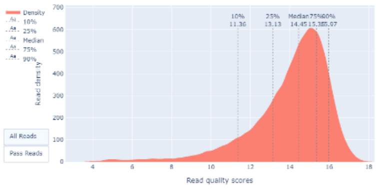


**Supplementary Figure3.**

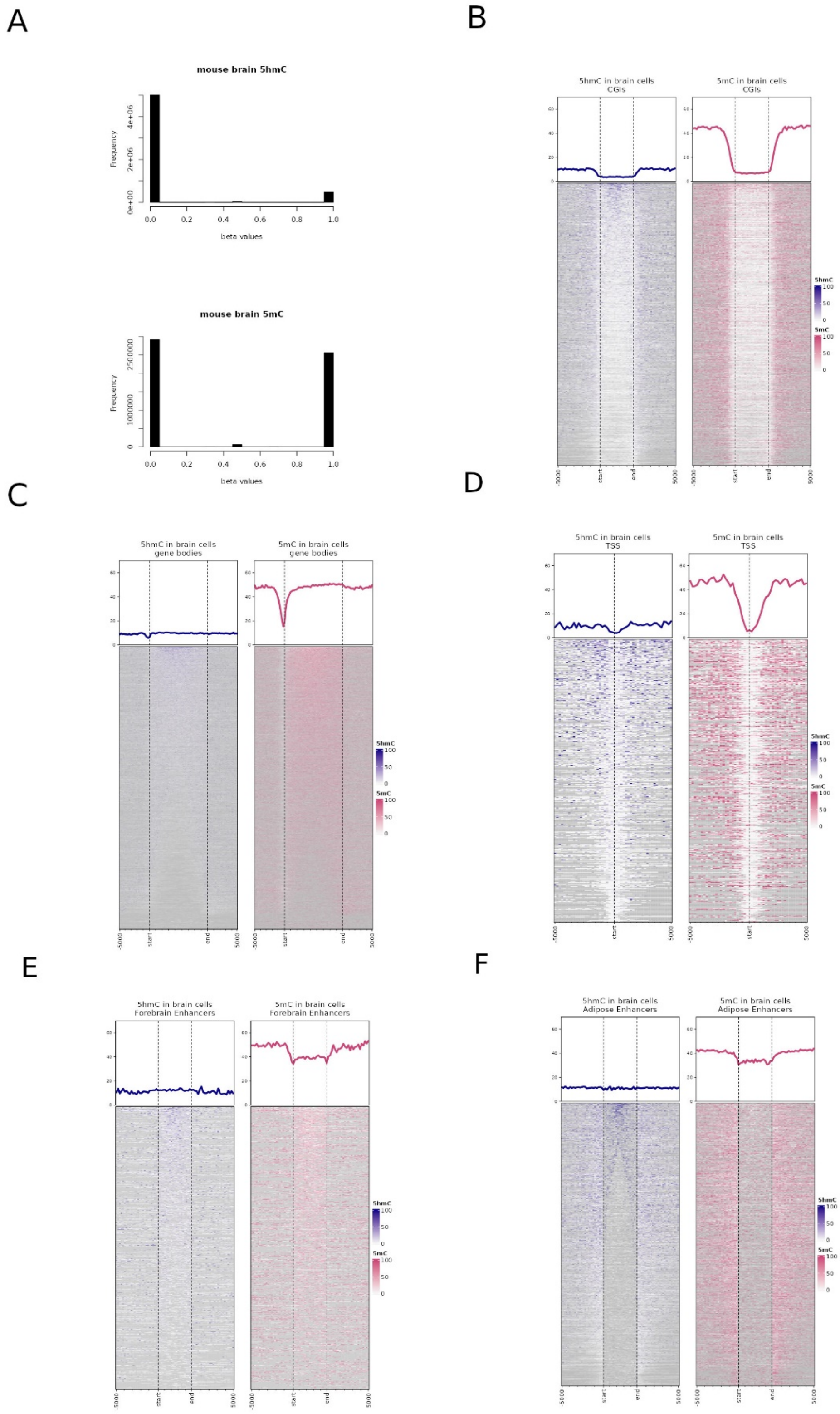


**Supplementary Figure 4.**

A

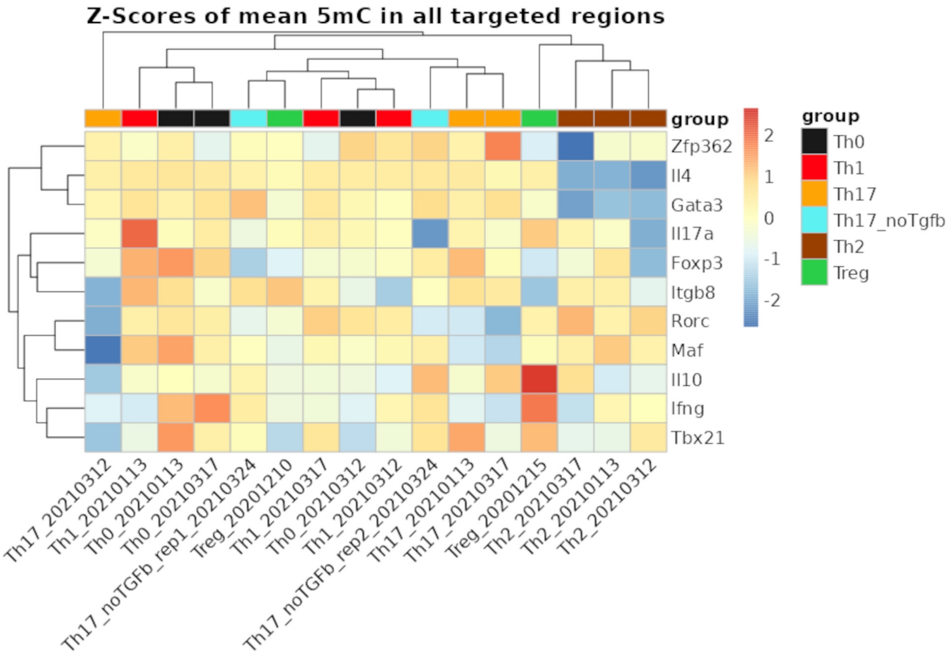


B

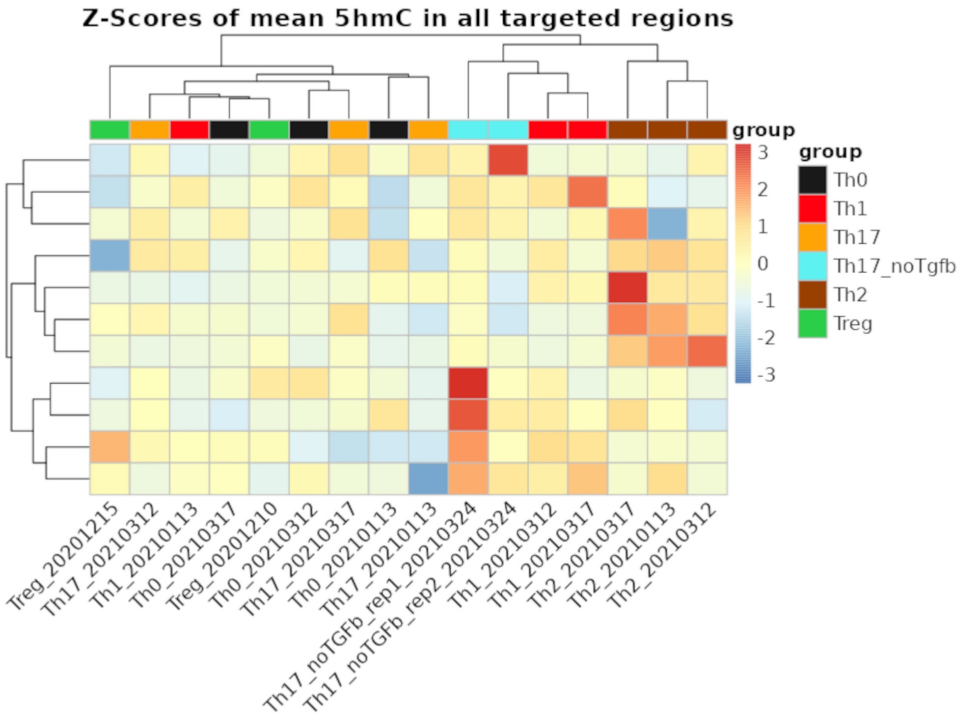


**Supplementary Figure 5**.

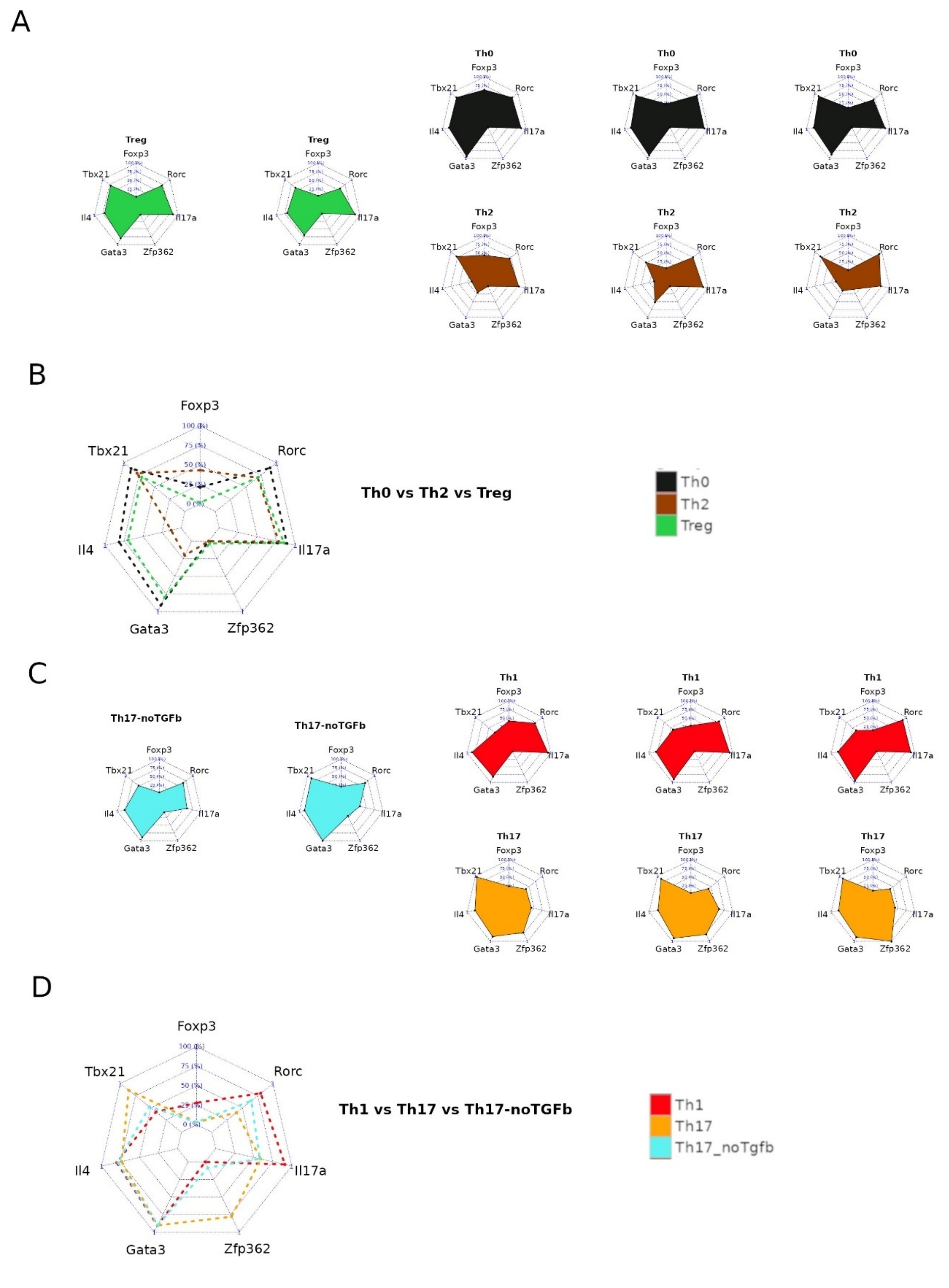


**Supplementary Figure 6**.

### Supplementary Tables

#### Supplementary Table 1.

| **Antibodies, recombinant proteins and Flow cytometry reagents** | | |
|---|---|---|
| Anti-mTbet- PE (clone 4B10) | eBioscience/ThermoFisher | Cat# 12-5825-82 |
| Anti-Rorɣt- BV650 (clone Q31-378) | BD Biosciences | Cat# 564722 |
| Anti-mFoxp3- PC7 (clone FJK-16S) | eBioscience/ThermoFisher | Cat# 25-5773-82 |
| Anti-GATA3-A647 (clone L50-823) | BD Biosciences | Cat# 560068 |
| CBA Flex Set mIFNγ | BD Biosciences | Cat#558296 |
| BD CBA Flex Set mIL-4 | BD Biosciences | Cat#558298 |
| BD CBA Flex Set mIL-17A | BD Biosciences | Cat#560283 |
| BD CBA Flex Set mIL-10 | BD Biosciences | Cat#558300 |
| Mouse/Rat Soluble Protein Master Buffer Kit | BD Biosciences | Cat#558266 |
| Mouse/Rat Soluble Protein Master Buffer Kit | BD Biosciences | Cat#558266 |
| Fix/perm kit : eBioscience FOXP3 /Transcription | eBioscience/ThermoFisher | Cat# 00-5523-00 |
| Live/Dead v506 | eBioscience/ThermoFisher | Cat#65-0866-14 |
| Live/Dead eF780 | eBioscience/ThermoFisher | Cat#65-0865-18 |
| InVivoMAb anti-mouse IL-4 (clone 11B11) | BioXcell | Cat# BE0045 |
| InVivoMAb anti-mouse CD3ε (clone 145-2C11) | BioXcell | Cat# BE0001-1 |
| InVivoMAb anti-mouse CD28 (clone PV1) | BioXcell | Cat# BE0015-5 |
| ProLeukin IL-2 « Clinical grade reagent » | N/A | N/A |
| Human recombinant TGF-β1 | R&D Systems | Cat#240-B |
| Murine recombinant IL12 | R&D Systems | Cat#419-ML |
| Murine recombinant IL4 | R&D Systems | Cat#404-ML |
| Murine recombinant IL6 | R&D Systems | Cat#406-ML |
| Murine recombinant IL1b | R&D Systems | Cat#401-ML |

**Supplementary Table 2**.

| **Cells** | **Polarizing condirions** |
|---|---|
| Th1 | IL-12 (10-15 ng/ml) and α-IL-4 antibody (5-10 µg/ml) |
| Th2 | IL-4 (15 ng/ml) and α-Ifnɣ antibody (10 µg/ml) |
| Th17 | IL-1b (10-20 ng/ml), IL-6 (20-40 ng/ml), hTGF-b (1-3 ng/ml) and α-IL-4 antibody (5-10 µg/ml), α-Ifnɣ antibody (10 µg/ml) |
| Th17- TGF-β | IL-1b (20 ng/ml), IL-6 (40 ng/ml), and α-IL-4 anti-body (5 µg/ml), α-Ifnɣ antibody (10 µg/ml) |
| Treg | hTGF-b (2,25-3 ng/ml) and rhIL-2 (100 U/ml). |

**Supplementary Table 3**. sgRNAs

| Gene Symbol | Strand | Sequence | PAM | On-T Score | Off-T Score |
|---|---|---|---|---|---|
| FoxP3_down2.1 | - | ACGGTGGAATTGCTGCCTGA | TGG | 57 | 68 |
| FoxP3_down2.2 | - | ATGGACTGCCCTGATAGATA | GGG | 62 | 57 |
| Foxp3_up_3.1 | - | TCGTCCGCACTCCTCATCCT | TGG | 51 | 71 |
| Foxp3_up_3.2 | - | ATGAGAGCCCTACGCAATCA | TGG | 58 | 81 |
| GATA3 down | - | GTTAGTTGTACACGGTACTT | CGG | 65 | 86 |
| GATA3 Up | + | AAGCTTGTAGTACAGCCCAC | AGG | 77 | 67 |
| Gata3_downX_2.2 | + | AGAGACCATAACAATAACGC | GGG | 68 | 71 |
| Gata3_upX_2.2 | - | ATTAGCGTTCCTCCTCCAGA | GGG | 53 | 65 |
| Ifnɣ_down | - | CAATGCCTTTCCAAGGGTAT | TGG | 71 | 50 |
| Ifnɣ_down_2 | - | GTCTTGCCTGGAATCCAAAT | GGG | 69 | 50 |
| Ifnɣ_up | + | GCATCTGGGTCAAGATAACT | GGG | 75 | 61 |
| Ifnɣ_up_2 | + | AATTGAGCCACTAGGAATGC | CGG | 73 | 60 |
| il10_down | - | GCAAGCCTGACATTGACGTG | CGG | 64 | 77 |
| il10_down_2 | - | TCTGACTAGTTATCATGTGC | TGG | 65 | 47 |
| il10_up | + | GACCTCACATAAGGTTCTTG | AGG | 74 | 68 |
| il10_up_2 | + | AAGGGCAACTAGCTGACAAA | TGG | 80 | 40 |
| il17a_down | - | TGTGGAACCTAAACACACGA | GGG | 82 | 73 |
| il17a_down_2 | - | ATGTGGAACCTAAACACACG | AGG | 80 | 64 |
| il17a_up | + | CTAGCTTTACCAATTCCATA | AGG | 75 | 46 |
| il17a_up_2 | + | ATTCCATAAGGCCTCCCATG | TGG | 64 | 43 |
| IL-4 Down | - | GGGGCAATGAGTACCTCGAC | AGG | 73 | 88 |
| IL-4 Up | + | GTTCTTGTTTCACAAGCCGC | AGG | 52 | 72 |
| il4_downX_2.2 | + | ATAGGTAAAGCCTCATTCCA | TGG | 71 | 58 |
| il4_upX_2.2 | - | TACCTCTGGATTCATCCCCC | TGG | 56 | 50 |
| MAF Down | - | CCATTTGAGCCTGACGTCAC | GGG | 77 | 46 |
| MAF UP | + | GGAAAGCTATCACACCTGTT | TGG | 78 | 50 |
| maf_downX_2.2 | + | TGGTGCACTAAACTTCGCAC | CGG | 53 | 85 |
| maf_upX_2.1 | - | TGATACATGGCTAAGTGCAG | AGG | 76 | 32 |
| rorc_down | - | AAGACCTAACTACCTAGCAC | AGG | 85 | 44 |
| rorc_down_2 | - | CAGGACACGACTGTTAAAAC | TGG | 61 | 71 |
| rorc_up | + | GATAAGAGGACTGGGCACGT | GGG | 76 | 71 |
| rorc_up_2 | + | GTCACGTTATGAGGTGCTGT | AGG | 64 | 73 |
| tbx21_downX_2.1 | + | CCCTACGGGTGAAGTCCTAT | TGG | 63 | 92 |
| tbx21_downX_2.2 | + | CCTACGGGTGAAGTCCTATT | GGG | 56 | 90 |
| tbx21_upX_2.1 | - | TCTCCAACCAATCACTATAC | AGG | 87 | 66 |
| tbx21_upX_2.2 | - | GCTGTCGCCACTGGAAGGAT | AGG | 64 | 68 |
| zfp362_downX_2 | + | GCTAAACTGGAATCCTCACG | AGG | 70 | 75 |
| zfp362_upX_2 | - | GCCAACCTTGGTCAACTCGG | TGG | 65 | 84 |

**Supplementary Tables 4 to 7 can be downloaded from the GitHub repository:**

*https://github.com/hernandezvargash/Tcell.5mC.ID*
